# Mechanistic insight into the interactions of NAP1 with NDP52 and TAX1BP1 for the recruitment of TBK1

**DOI:** 10.1101/264085

**Authors:** Tao Fu, Jianping Liu, Yingli Wang, Xingqiao Xie, Shichen Hu, Lifeng Pan

**Affiliations:** State Key Laboratory of Bioorganic and Natural Products Chemistry, Center for Excellence in Molecular Synthesis, University of Chinese Academy of Sciences, Shanghai Institute of Organic Chemistry, Chinese Academy of Sciences, 345 Lingling Road, Shanghai 200032, China.

**Keywords:** Autophagy receptor, NDP52, NAP1, TAX1BP1, TBK1, Selective autophagy

## Abstract

NDP52 and TAX1BP1, two SKICH domain-containing autophagy recetpors, play crucial roles in selective autophagy. The autophagic functions of NDP52 and TAX1BP1 are regulated by TBK1, which can indirectly associate with them through the adaptor protein NAP1. However, the molecular mechanism governing the interactions of NAP1 with NDP52 and TAX1BP1 as well as the effects induced by TBK1-mediated phosphorylation of NDP52 and TAX1BP1 remain elusive. Here, we reported the first atomic structures of the SKICH regions of NDP52 and TAX1BP1 in complex with NAP1, which not only uncover the mechanismtic basis underpinning the specific interactions of NAP1 with NDP52 and TAX1BP1, but also reveal the first binding mode of a SKICH domain. Moreover, we demonstrated that the phosphorylation of TAX1BP1 SKICH mediated by TBK1 may regulate the interaction between TAX1BP1 and NAP1. In all, our findings provide mechanistic insights into the NAP1-mediated recruitments of TBK1 to NDP52 and TAX1BP1, and are valuable for further understanding the functions of these proteins in selective autophagy.

## Introduction

Autophagy is a tightly regulated lysosome-dependent “self-eating” catabolic process to recycle cytoplasmic material in eukaryotic cells, and plays a critical role in the maintenance of cellular homeostasis and physiology (Jiang & Mizushima, 2014; Klionsky & Emr, 2000; Mizushima, 2007). Previously, autophagy was regarded as a non-specific bulk degradation process with little or no selectivity. Recently, increasing evidences reveal that many cytosolic contents, including bulk protein aggregates, dysfunctional organelles and invading pathogens, are degraded by autophagy in a highly selective manner, which is termed as selective autophagy (Feng et al, 2014; Gomes & Dikic, 2014; Mizushima, 2007; Nakatogawa et al, 2009; Randow & Youle, 2014; Stolz et al, 2014). The key factors involved in selective autophagy processes are autophagy receptors such as SQSTM1/p62, NDP52, TAX1BP1, Optineurin, NBR1, Nix, FUNDC1, FAM134B, and NCOA4 in mammals, which not only can specifically recognize relevant autophagic cargoes but also can bind to the key autophagic factor, the ATG8 family proteins, thereby serving as bridging adaptors to target specific cargoes to the autophagy machinery for subsequent autophagic degradations (Johansen & Lamark, 2011; Rogov et al, 2014; Stolz et al, 2014). In view of crucial roles played by autophagy receptors in selective autophagy, the autophagic functions of autophagy receptors have been well spatially and temporally tuned by other regulatory proteins, particularly protein kinases, such as the Casein kinase 2 (CK2) and the TANK-binding kinase 1 (TBK1) (Heo et al, 2015; Matsumoto et al, 2011; Richter et al, 2016; Wild et al, 2011). For instance, TBK1 was found to phosphorylate the Ser177 residue in Optineurin and the Ser403 residue in SQSTM1 to promote their interactions with ATG8 family proteins and ubiquitin proteins, respectively (Matsumoto et al, 2011; Wild et al, 2011). However, until now, many of the detailed molecular mechanism underpinning the specific associations of autophagy receptors with their regulatory proteins as well as the down-steam effects mediated by these regulatory proteins in selective autophagy processes are still elusive.

NDP52 and TAX1BP1 are two important ubiquitin-binding and multi-domain autophagy receptors in mammals, both of which are demonstrated to participate in selective autophagic degradations of invading infectious pathogens (xenophagy) such as *Salmonella enterica Typhimurium*, and the depolarized mitochondria (mitophagy) (Lazarou et al, 2015; Newman et al, 2012; Thurston et al, 2009; Tumbarello et al, 2015). In addition, NDP52 was also reported to mediate selective autophagic degradations of retrotransposon RNA (Guo et al, 2014), and specific functional proteins including DICER, AGO2 in the miRNA pathway and MAVS in the immune signaling (Gibbings et al, 2012; Jin et al, 2017). Notably, genetic mutation of NDP52 is directly linked with Crohn’s disease, a type of inflammatory bowel disease likely caused by a combination of environmental, immune, and bacterial factors (Ellinghaus et al, 2013). Except for the diverse central coiled-coil region, NDP52 and TAX1BP1 share a highly similar domain structure (Fig 1A), and both contain a N-terminal SKIP carboxyl homology (SKICH) domain followed by an unconventional LC3-interacting region (LIR) motif that can differentially bind to different LC3/GABARAP orthologues (Tumbarello et al, 2015; von Muhlinen et al, 2012), and two C-terminal zinc fingers (Fig 1A), which participate in the recognitions of ubiquitin proteins decorated on autophagic cargoes and the unconventional myosin motor Myosin VI (Tumbarello et al, 2015; Tumbarello et al, 2012; Xie et al, 2015). In contrast to TAX1BP1, NDP52 uniquely contains a Galectin8 interacting region (GIR) (Fig 1A), which can specifically interact with the sugar receptor Galectin8 to target vesicle-damaging pathogens (Kim et al, 2013; Li et al, 2013; Thurston et al, 2012). In addition to NDP52 and TAX1BP1, SKICH domains are also found in the CALCOCO1, SKIP and proline rich inositol-polyphosphate 5-phosphatase (PIPP) proteins, the SKICH domains of which are important for plasma membrane localization (Gurung et al, 2003). However, due to the lack of systemic characterization, the precise working mode of SKICH domain is still not well established. Interestingly, the N-terminal SKICH domains of NDP52 and TAX1BP1 are reported to interact with the adaptor protein NAP1 and SINTBAD (Thurston et al, 2009), both of which can further directly bind to the TBK1 kinase through their C-terminal TBK1-binding domains (TBD) (Li et al, 2016; Ryzhakov & Randow, 2007) (Fig 1A). Therefore, NDP52 and TAX1BP1 can indirectly associate with TBK1 through NAP1 or SINTBAD. Importantly, recent studies reveal that the recruitment of TBK1 as well as its kinase activity are required for efficient xenophagy and mitophagy processes (Heo et al, 2015; Lazarou et al, 2015; Moore & Holzbaur, 2016; Richter et al, 2016; Thurston et al, 2016). Strikingly, TBK1 can directly mediate the phosphorylation of NDP52 and TAX1BP1 at multiple sites including their SKICH domains (Heo et al, 2015; Richter et al, 2016). However, how TBK1 associates with NDP52 and TAX1BP1 mediated by NAP1 or SINTBAD as well as the down-steam consequences induced by TBK1-mediated phosphorylation of NDP52 and TAX1BP1 are currently unknown, and the detailed binding mechanism of NDP52 and TAX1BP1 with the TBK1-binding adaptors remains to be elucidated.

**Figure 1.**
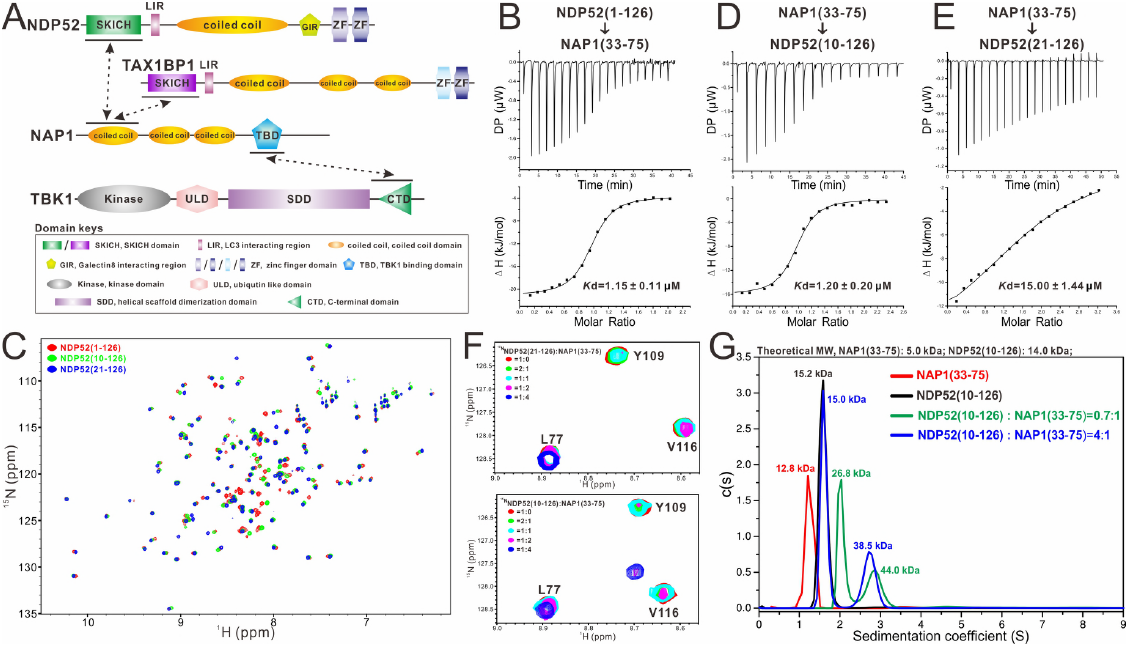
Biochemical characterizations of the interaction between NDP52 and NAP1. (**A**) A schematic diagram showing the domain organizations of NDP52, TAX1BP1, NAP1 and TBK1. In this drawing, the NDP52/NAP1, TAX1BP1/NAP1 and NAP1/TBK1 interactions are further highlighted and indicated by two-way arrows. (**B)** The ITC-based measurement of the binding affinity of NAP1(33-75) with NDP52(1-126). The *K*d error is the fitted error obtained from the data analysis software, when using the one-site binding model to fit the ITC data. (**C**) Overlay plot of the ^1^H-^15^N HSQC spectra of three NDP52 fragments, NDP52(1-126), NDP52(10-126), and NDP52(21-126). (**D** and **E**) ITC-based measurements of the binding affinities of NAP1(33-75) with NDP52(10-126) (**D**), and NDP52(21-126) (**E**). The *K*d errors are the fitted errors obtained from the data analysis software, when using the one-site binding model to fit the ITC data. (**F**) Superposition plots of two selected regions of the ^1^H-^15^N HSQC spectra of NDP52(10-126) and NDP52(21-126) titrated with increasing molar ratios of the NAP1(33-75) proteins. (**G**) Overlay plot of the sedimentation velocity data of NAP1(33-75), NDP52(10-126), and NAP1(33-75) incubated with 1:0.7 or 1:4 molar ratio of NDP52(10-126). The results demonstrate that NAP1(33-75) form a stable dimer and may interact with one or two monomeric NDP52(10-126) to form a 2:1 or 2:2 stoichiometric complex in solution.

In this study, we biochemically and structurally characterized the interactions between NAP1 and the two autophagy receptors, NDP52 and TAX1BP1, and discovered that the N-terminal coiled-coil domain of NAP1 forms a stable dimer and specifically interact with the SKICH regions of NDP52 and TAX1BP1 to form two unique hetero-tetramers. The determined structures of NDP52/NAP1 and TAX1BP1/NAP1 complexes not only uncovered the detailed binding mechanism of NAP1 with NDP52 and TAX1BP1, but also unveiled that NDP52 and TAX1BP1 share a general binding mode to interact with NAP1 and SINTBAD. Furthermore, we evaluated the currently known phosphorylation sites mediated by TBK1 in the SKICH domains of NDP52 and TAX1BP1, and discovered that the phosphorylation of TAX1BP1 S25 residue may regulate the interaction between TAX1BP1 and NAP1. In summary, our findings provided mechanistic insights into the interactions of NAP1 with autophagy receptor NDP52 and TAX1BP1 for the recruitment of TBK1, and expanded our understandings of the functions as well as the working modes of NDP52, TAX1BP1, NAP1 and TBK1 in selective autophagy.

## Results

### Biochemical characterizations of the specific interaction between NDP52 and NAP1

To gain molecular insights into the specific interaction of NDP52 with NAP1, we first conducted a detailed sequence alignment analysis of the NDP52 SKICH domain and the NAP1 N-terminal coiled-coil regions, which were reported to mediate the NDP52/NAP1 complex formation in an earlier study (Thurston et al, 2009). The result showed that these regions of NDP52 and NAP1 are highly conserved during evolution (**Fig EV1**), in line with their potential functional roles to interact with each other. To further validate and define the precise binding regions of NDP52 and NAP1, we carried out a quantitative isothermal titration calorimetry (ITC)-based assay using the entire NDP52 SKICH region (residues 1-126) and three different NAP1 fragments, NAP1(1-85), NAP1(1-75) and NAP1(33-75). Our result demonstrated that the three NAP1 fragments bind to NDP52(1-126) with similar *K*_D_ values, about 1.49 μM, 1.80 μM and 1.15 μM, respectively (**Figs EV2A, B** and **1B**), indicating that the NDP52 SKICH region indeed can directly bind to the NAP1 N-terminal coiled-coil region, and the NDP52-binding site of NAP1 is located within the NAP1(33-75) fragment. Unfortunately, our initial attempts to obtain protein crystals using the purified NDP52(1-126)/NAP1(33-75) complex failed, therefore we sought to further narrow down the NAP1-binding region of NDP52 in order to get a more suitable protein complex for subsequent structural characterizations.

Based on sequence conservation and secondary structure prediction of NDP52 (**Fig EV1B)**, we constructed two additional NDP52 fragments, NDP52(10-126) and NDP52(21-126), which lack different extreme N-terminal loop residues in compare with the NDP52(1-126) fragment. Further NMR-based analysis showed that the well-dispersed ^1^H-^15^N HSQC spectra of those three NDP52 fragments are very similar (Fig 1B), suggesting that those three NDP52 proteins are well-folded and adopt a similar overall structure. However, in contrast to NDP52(10-126), which binds to NAP1(33-75) with an affinity comparable to that of NDP52(1-126), the NDP52(21-126) fragment displays a much weaker binding affinity towards NAP1(33-75) as indicated by our ITC-based analysis (Fig 1B, D and E). Thus, the NDP52(10-126) fragment rather than the NDP52(21-126) fragment is sufficient for binding to NAP1. To further confirm this notion, we also used NMR spectroscopy to characterize the interactions of NAP1(33-75) with NDP52(10-126) and NDP52(21-126). Titrations of ^15^N-labeled NDP52(10-126) and NDP52(21-126) with un-labeled NAP1(33-75) proteins showed that a select set of peaks in the ^1^H-^15^N HSQC spectra of NDP52(10-126) and NDP52(21-126) underwent significant peak-broadenings or chemical shift changes (**Fig EV2C** and **D**), indicating that those two NDP52 proteins can directly bind to NAP1(33-75). However, further detailed analyses revealed that many peaks in the ^1^H-^15^N HSQC spectra of these two proteins (**Fig EV 2C** and **D**), such as the peak corresponding to the V116 residue, showed very different change profiles when titrated with NAP1(33-75), and importantly, a slow exchange pattern was observed in the NMR titration experiment of NDP52(10-126) (Fig 1F), confirming that the NDP52(10-126) and NDP52(21-126) fragments differentially bind to NAP1(33-75), and NDP52(10-126) interacts with NAP1(33-75) more strongly than the NDP52(21-126) fragment. Furthermore, using an analytical ultracentrifugation-based assay, we revealed that NAP1(33-75) forms a stable homo-dimer, and NDP52(10-126) is a monomer in solution (Fig 1G). Interestingly, although NAP1(33-75) can specifically interact with NDP52(10-126) to form a single complex peak on an analytical gel filtration chromatography analysis (**Fig EV2E**), our sedimentation velocity results revealed that the NAP1(33-75) dimer actually may bind to one or two NDP52(10-126) molecules to form a 1:1 or 1:2 stoichiometric complex under a un-saturated condition (Fig 1G). However, in the presence of excess amounts of NDP52(10-126) proteins, the NAP1(33-75) dimer can simultaneously bind to two NDP52(10-126) molecules to form a hetero-tetramer (Fig 1G).

### Overall structures of NDP52(10-126) and its complex with NAP1(33-75)

To further uncover the mechanistic basis underlying this unique interaction between NDP52(10-126) and NAP1(33-75), we sought to determine their atomic structures. Although the structure of NDP52(21-141) fragment was solved in a previous study(von Muhlinen et al, 2012), based on our aforementioned ITC and NMR results, the NDP52(10-126) fragment is different from NDP52(21-126) in binding to NAP1. Therefore, we first determined the crystal structure of NDP52(10-126) using the molecular replacement method (**Table EV1**). Notably, in the crystal structure, the extreme N-terminal 8 residues of NDP52(10-126) are unsolved due to lacking of electron density, but the critical F20 residue of NDP52 for interacting with NAP1 (see below for more details) is well defined. The structure of NDP52(10-126) features a β-sandwich Ig-like architecture assembled by a 3-stranded antiparallel β-sheet packing with a 4-stranded antiparallel β-sheet (**Fig EV3A**). Unsurprisingly, the overall structure of NDP52(10-126) is highly similar to the previously reported structure of NDP52(21-141) except for the N-terminal and C-terminal loop regions (**Fig EV3B**).

Next, we purified the NDP52(10-126)/NAP1(33-75) complex in the presence of excess amounts of NDP52(10-126), and successfully obtained good crystals that diffracted to 2.02 Å resolution. Using the molecular replacement method with the apo-form structure of NDP52(10-126), we managed to solve the complex structure (**Table EV1**). In the final complex structural model, each asymmetric unit contains one NDP52(10-126)/NAP1(33-75) complex, which has a 2:2 stoichiometry and forms a symmetrical hetero-tetramer consisting of two NDP52 molecules and a NAP dimer (Fig 2A), consistent with our analytical ultracentrifugation analysis (Fig 1G). In the complex structure, the two NAP1 molecules mainly form two continuous α-helices, which head-to-head pack against each other to form a parallel but crossed coiled-coil homo-dimer (Fig 2), and intriguingly, the two NDP52 molecules through each solvent-exposed side of the 4-stranded β-sheet symmetrically bind to the N-terminal region of the NAP1 dimer, forming a unique hetero-tetrameric complex with an overall architecture distinct from any currently known protein structures revealed by a structural similarity search using the program Dali (Holm & Rosenstrom, 2010) (Fig 2A). Notably, there is no direct contact between two NDP52 molecules in the complex structure (Fig 2). Further structural comparison revealed that the NDP52(10-126) molecules in the complex structure also miss their N-terminal 8 residues owing to the lack of electron density, and adopts a similar overall conformation to that of the apo-form protein, except for the extreme N-terminal loop that is directly involved in the NAP1-binding, and the flexible loop linking β4 and β5 (**Fig EV3C**).

**Figure 2.**
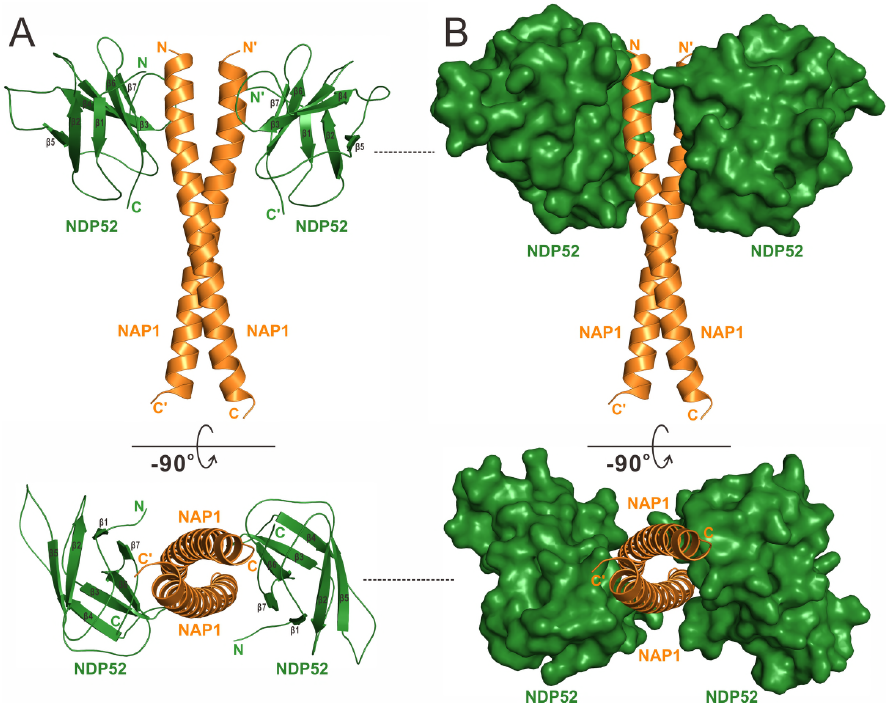
The overall structure of NDP52(10-126) in complex with NAP1(33-75). **(A)** Ribbon diagram showing the overall structure of NDP52(10-126) in complex with NAP1(33-75). In this drawing, NDP52(10-126) is shown in forest green, NAP1(33-75) in orange. **(B)** The combined ribbon and surface representation showing the overall architecture of NDP52/NAP1 complex with the same color scheme as in panel A.

### The dimerization interface of NAP1 in the NDP52/NAP1 complex

Further structural analysis revealed that in the structure of NDP52(10-126)/NAP1(33-75) complex, the coiled-coil region (residues 36-73) of NAP1 is composed of 5 regular heptad repeats, and is responsible for the NAP1 dimer formation (Figs 2 and 3A-C). The dimerization of NAP1 is mediated by extensive hydrophobic and polar interactions between residues located at the *b*, *c*, *f*, *g* positions of the two NAP1 coiled-coil helices (Fig 3A-C). Particularly, a N-terminal hydrophobic patch formed by L41, V42, A44, Y45, I48, K49, and L52 residues together with a C-terminal hydrophobic patch assembled by L62, K63, I66, L69, and L73 residues of one NAP1 helix pack against their corresponding counterparts in the other NAP1 helix to form two separated hydrophobic interfaces of the NAP1 dimer (Fig 3A-C). Strikingly, a highly specific polar interaction network comprised by un-symmetrical hydrogen bonds and salt bridges was found between the side chains of R51, S55, E56, E58, N59, and K63 residues located in the middle region of the paired NAP1 helixes (Fig 3A, B, and D). Moreover, two symmetrical inter-molecular hydrogen bonds formed between the side chains of S37 in the *b* position of one NAP1 helix and H38 in the *c* position of the other NAP1 helix as well as two charge-charge interactions found between the positively charged R65 residue of one NAP1 helix and the negatively charged E70 of the other NAP1 helix further stabilize the NAP1 dimer formation (Fig 3A-C).

**Figure 3.**
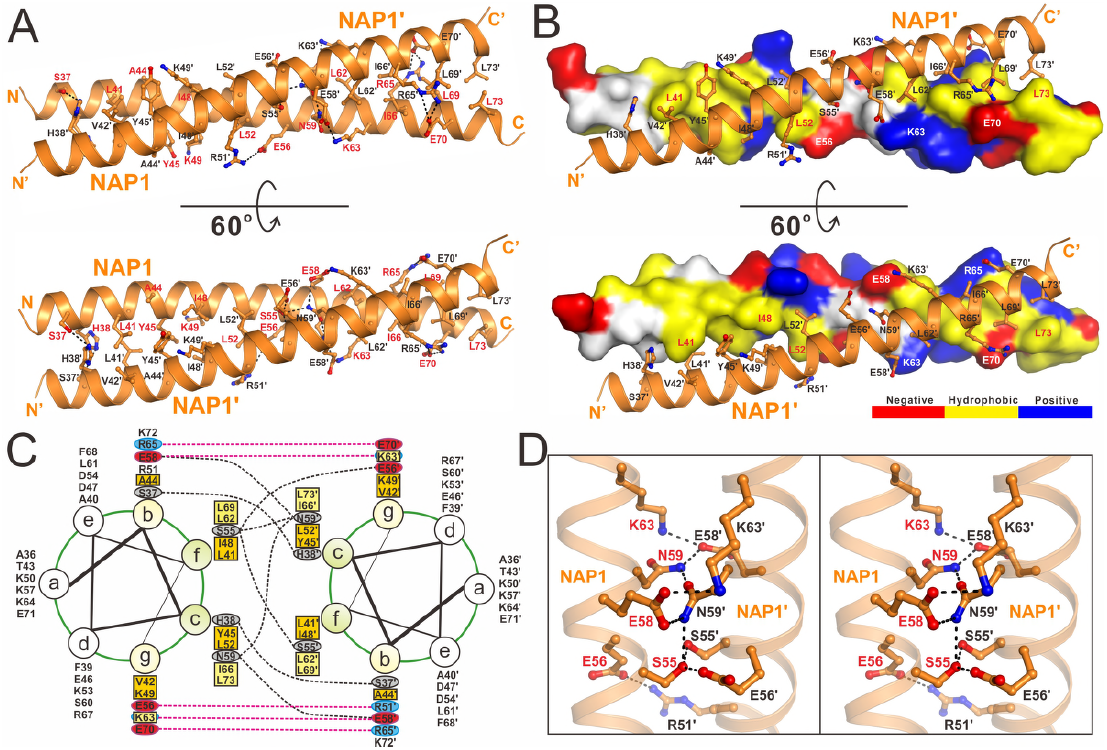
The dimerization interface of NAP1 dimer in the NDP52/NAP1 complex. **(A)** The combined ribbon and stick-ball representation showing the detailed interactions between two NAP1 monomers in the NDP52/NAP1 complex. In this drawing, the side chains of the key residues are shown in the stick-ball mode, and the hydrogen bonds involved in the binding are shown as dotted lines. **(B)** The combined surface representation and the ribbon-stick model showing the molecular interface of NAP1 dimer in the NDP52/NAP1 complex. In this presentation, one NAP1 monomer is shown in the surface model and the other in the ribbon-stick model. The hydrophobic amino acid residues of NAP1 monomer in the surface model are drawn in yellow, the positively charged residues in blue, the negatively charged residues in red, and the uncharged polar residues in gray. **(C)** A helical wheel presentation showing the detailed dimerization interface formed between the heptad repeats of two NAP1(33-75) monomers. In this presentation, the inter-helical hydrogen bonds and salt bridges are depicted by black and pink dashed lines, respectively. Two hydrophobic contacts located at the N-terminal and C-terminal parts of the NAP1 coiled-coil dimer and formed by hydrophobic residues located at the “*b*”, “*c*”, “*g*”, “*f*” positions are further highlighted with orange and yellow boxes, respectively. **(D)** Stereo view of the ribbon-stick representation showing the unique polar interaction network found in the middle region of the NAP1 dimerization interface. The relevant hydrogen bonds and salt bridges involved in the binding are shown as dotted lines.

### The binding interface between NDP52 and NAP1

In the complex, the two NDP52 molecules symmetrically bind to two homo-dimeric interfaces located at the opposite sides of the N-terminal part of the NAP1 dimer, each burying a total surface area of ∼824 Å^2^ (Fig 2). Detailed structural analysis showed that the binding interface between NDP52 and NAP1 is formed by residues from the β6, β7 and loops located at the solvent-exposed face of the 4-stranded β-sheet of NDP52, and accommodates NAP1 residues located at the N-terminal region of the paired helixes through both hydrophobic and polar interactions (Fig 4A). In particular, the hydrophobic side chains of V35, A36, F39 and A40 from one chain of NAP1 dimer pack against a hydrophobic patch formed by the side chains of F20, W63, C108, V116 and A119 from NDP52, and the hydrophobic side chains of V61, P122 of NDP52 partially occupy a hydrophobic groove formed by the side chains of L41, A44, I48 from one chain of the NAP1 dimer and the aromatic side chain of Y45 from the other chain of the NAP1 dimer (Fig 4A). In addition, the backbone carboxyl groups of V61, G62, W63, K64 and P122 together with the polar side chain groups of Y104, Q106, Q124 from NDP52 interact with the S37, A40, R51 residues from one chain of the NAP1 dimer and the H38, Y45, E56 residues from the other chain of the NAP1 dimer to form nine highly specific hydrogen bonds (Fig 4A). Moreover, an Arg-Glu pair (Arg126_NDP52_-Glu56_NAP1_) and a Lys-Glu pair (Lys49_NAP1_-Glu103_NDP52_) of salt bridges further strengthen the NDP52 and NAP1 interaction (Fig 4A). In line with their important structural roles, all these key residues of NDP52 and NAP1 involved in the binding interface are evolutionarily conserved across different eukaryotic species (**Fig EV1**).

**Figure 4.**
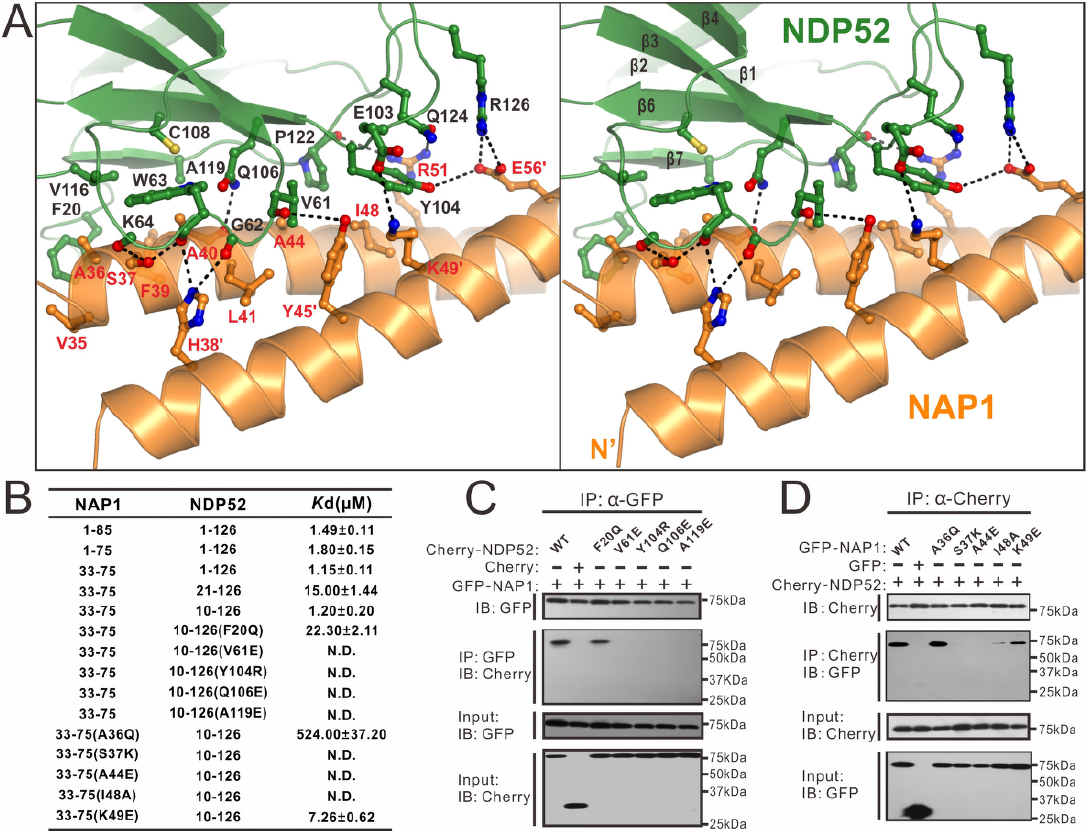
Molecular details of the NDP52 and NAP1 interaction. **(A)** Stereo view of the ribbon-stick representation showing the binding interface between NDP52 and NAP1 in the determined NDP52/NAP1 complex structure. The hydrogen bonds and salt bridges involved in the interaction are shown as dotted lines. **(B)** The measured binding affinities between various forms of NDP52 and NAP1 or their mutants by ITC-based assays. (**C** and **D**) Mutagenesis-based Co-IP assays to confirm the interactions between NDP52 and NAP observed in the complex structure.

Using ITC and Co-IP analyses, we further verified the specific interactions between NDP52 and NAP1 observed in the complex structure. Consistent with our structural data, the ITC results showed that the individual mutation of key interface residues either from NDP52 or NAP1, such as the V61E, Y104R, Q106E, A119E mutations of NDP52 or the A36Q, S37 K, A44E, I48A, K49E mutations of NAP1, essentially abolished or largely reduced the specific interaction between NDP52(10-126) and NAP1(33-75) (**Figs EV4** and **4B**). Notably, the replacement of F20 residue located at the N-terminal loop of NDP52 with a Gln residue reduced the binding affinity between NDP52(10-126) and NAP1(33-75) to a value comparable to that of NDP52(21-126) and NAP1(33-75) interaction (**Figs EV4A**, **1E** and **4B**), further confirming our aforementioned biochemical and structural results (**Figs 1C, D** and 4**A**). Importantly, in agreement with our *in vitro* ITC results, further co-immunoprecipitation experiments revealed that point mutations of key interface residues including the F20Q, V61E, Y104R, Q106E, A119E mutations of NDP52 and the S37 K, A44E, I48A, K49E mutations of NAP1 all completely disrupt or significantly attenuate the specific interaction between full-length NDP52 and NAP1 in co-transfected cells (Fig 4C and D). To our surprise, in contrast to the *in vitro* biochemical result, the A36Q mutation of NAP1 had no obvious effects on the interaction between NAP1 and NDP52 (Fig 4D), despite the exact reason for the observed discrepancy of NAP1 A36Q mutation on the interaction of NAP1 and NDP52 *in vitro* and *in vivo* remains to be elucidated. Nevertheless, all these data demonstrated that the specific interaction between NDP52(10-126) and NAP1(33-75) is essential for the NDP52/NAP1 complex formation.

### Cellular co-localization of NDP52 and NAP1 required the specific interaction between NDP52(10-126) and NAP1(33-75)

Next, we examined the role of NDP52(10-126)/NAP1(33-75) interaction on the cellular localizations of NDP52 and NAP1 in transfected HeLa cells. When co-transfected, the mCherry-tagged NDP52 displayed a punctate staining pattern, and co-localized perfectly with the GFP-tagged NAP1 puncta in the cytoplasm of transfected cells (Fig 5A and G). As control, the mCherry-tagged NDP52 and GFP-tagged NAP1 didn’t co-localize well with the GFP tag or the mCherry tag from the co-transfected empty vector (**Fig EV5**). In contrast, when co-transfection of NDP52 with the NAP1 S37 K or A44E mutant, each of which was demonstrated to lose its ability to interact with NDP52 (Fig 4B and D), the mCherry-tagged NDP52 became more diffused in cells, and the co-localization of NDP52 and NAP1 was largely compromised (Fig 5B, C and G). To further evaluate the NDP52(10-126)/NAP1(33-75) interaction in the cellular NAP1/NDP52 co-localization, we also assayed the V61E, Y104R and Q106E mutants of NDP52, as individual mutation of these three residues of NDP52 would essentially eliminate the NDP52/NAP1 interaction *in vitro* (Fig 4B and C). When the NDP52 V61E, Y104R or Q106E mutant was co-expressed with the wild type NAP1, it displayed a diffused localization pattern in the cytosol and its co-localization with NAP1 puncta was also dramatically reduced (Fig 5D-G), in line with our biochemical and structural results (Fig 4). Taken together, all these data clearly demonstrated that the specific interaction between NDP52(10-126) and NAP1(33-75) is critical for the cellular co-localization of NDP52 and NAP1.

**Figure 5.**
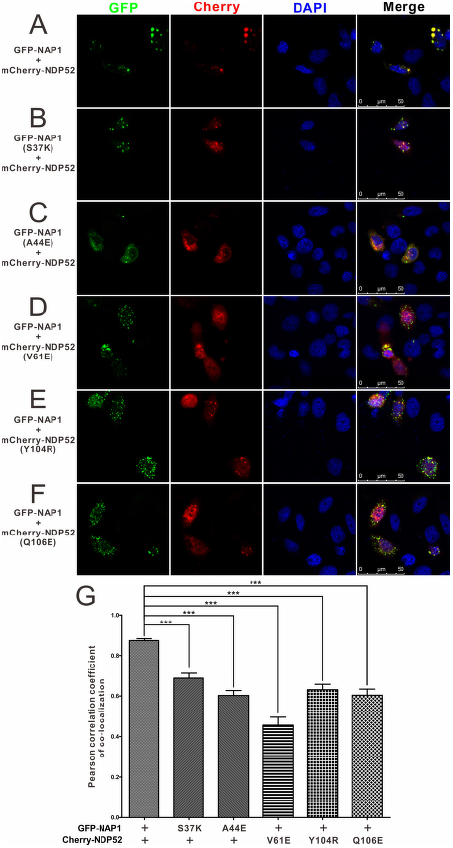
The specific interaction between NDP52(10-126) and NAP1(33-75) is required for the cellular co-localization of NDP52 and NAP1 in co-transfected HeLa cells. **(A)** When co-expressed, NDP52 co-localizes well with the NAP1 puncta. **(B-F)** Point mutations of key interface residues of NDP52 or NAP1 that disrupted their interaction *in vitro* largely decrease the co-localization of NDP52 and NAP1. **(G)** Statistical result related to the co-localizations of different NDP52 and NAP1 variants in co-transfected HeLa cells shown as Pearson correlation. The Pearson’s correlation coefficient analysis was performed using the LAS X software based on a randomly selected region that roughly contains one co-transfected HeLa cell. The data represent mean ± s.d. of >30 analyzed cells from two independent experiments. The unpaired Student t-test analysis was used to define a statistically significant difference, and the stars indicate the significant differences between the indicated bars (***P < 0.001).

### The molecular mechanism of TAX1BP1 and NAP1 interaction

In addition to autophagy receptor NDP52, NAP1 was also implicated in the binding to the SKICH-containing autophagy receptor TAX1BP1 (Thurston et al, 2009). However, the detailed binding mechanism remains elusive. Therefore, we also decided to characterize the interaction between NAP1 and TAX1BP1. Careful sequence alignment analysis showed that the SKICH region of TAX1BP1 (residues 1-121) is highly conserved during the evolution (**Fig EV6A**), and is very similar to that of NDP52 (Fig 6A). Analytical gel filtration chromatography and ITC-based assays revealed that the SKICH region of TAX1BP1 can directly interact with NAP(33-75) with a binding affinity *K*_D_ value of ∼2.05 μM (**Fig EV6B** and **C**), which is close to the *K*_D_ value of the NDP52/NAP1 interaction (Fig 1C).

**Figure 6.**
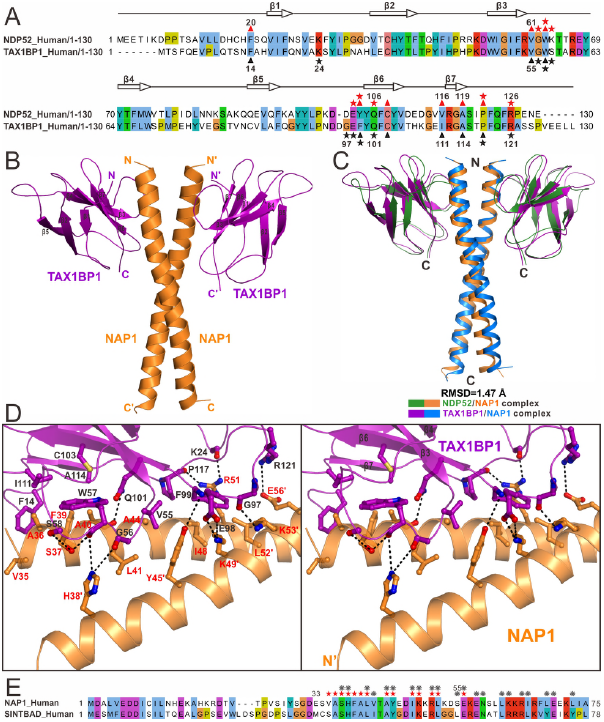
TAX1BP1 and NDP52 share a general binding mode to interact with NAP1 and SINTBAD. **(A)** The structure-based sequence alignment analysis of the SKICH regions of NDP52 and TAX1BP1. In this alignment, the conserved residues are highlighted by colors using software Jalview2.8.1 (http://www.jalview.org). The key residues of NDP52 that interact with NAP1 are highlighted with red stars (polar interaction) or red triangles (hydrophobic interaction), and that of TAX1BP1 in black. **(B)** Ribbon diagram showing the overall structure of TAX1BP1(1-121) in complex with NAP1(33-75). **(C)** Structural comparison of the overall structures of the TAX1BP1(1-121)/NAP1(33-75) complex and the NDP52(10-126)/NAP1(33-75) complex. In this drawing, the TAX1BP1(1-121) and NAP1(33-75) molecules in the TAX1BP1(1-121)/NAP1(33-75) complex are shown in purple and in blue, respectively, while the NDP52(10-126) and NAP1(33-75) in the NDP52(10-126)/NAP1(33-75) complex are drawn in green and in orange, respectively. **(D)** Stereo view of the ribbon-stick representation showing the detailed binding interface of the TAX1BP1/NAP1 complex. The related hydrogen bonds and salt bridges involved in the TAX1BP1 and NAP1 interaction are shown as dotted lines. **(E)** Sequence alignment analysis of the NDP52-binding regions in NAP1 and SINTBAD showing that most key interface residues of NAP1 that are crucial for dimerization or interacting with NDP52 are also conserved in SINTBAD. The interface residues of NAP1, which are critical for the interactions with NDP52 and/or TAX1BP1, are highlighted with red stars, while the NAP1 residues that are involved in the dimerization of NAP1 are labeled with grey gears.

To uncover the detailed binding mode between TAX1BP1 and NAP1, we also determined the high resolution crystal structure of the TAX1BP1(1-121)/NAP1(33-75) complex (**Table EV1**). As expected, the TAX1BP1(1-121)/NAP1(33-75) complex structure is composed of a NAP1 coiled-coil dimer and two TAX1BP1 molecules, which pack together to form a symmetrical hetero-tetramer (Fig 6B). The overall conformations of TAX1BP1(1-121)/NAP1(33-75) complex as well as the NAP1 dimer in the TAX1BP1/NAP1 complex are very similar to that of the NDP52/NAP1 complex (Figs 6C and **EV7A**). In the complex structure, the TAX1BP1(1-121) molecule misses the 11 residues located at the N-terminal loop owing to the lack of electron density, and adopts a similar fold to that of NDP52(10-126) in the NDP52/NAP1 complex (**Fig EV7B**).

Further structural analysis revealed that the association between TAX1BP1(1-121) and NAP1(33-75) is driven by extensive hydrophobic contacts and polar interactions (Figs 6D, **EV7C** and **D**). Specifically, the hydrophobic side chains of V35, A36, F39 and A40 from one chain of the NAP1 dimer contact extensively with the hydrophobic side chains of F14, W57, C103, I111 and A114 from TAX1BP1 (Figs 6D and **EV7C**), and the hydrophobic side chains of TAX1BP1 V55, F99, and P117 fit into a hydrophobic pocket assembled by the side chains of L41, A44, I48 residues from one chain of the NAP1 dimer and Y45, K49, L52 from the other chain (Fig 6D). Interestingly, similar to the corresponding interaction found in the NDP52/NAP1 complex (Fig 4A), a unique polar Q101 residue of TAX1BP1 is buried in the hydrophobic core of the interface by forming a strong hydrogen bond with the backbone carboxyl group of NAP1 A40 residue (Fig 6D). In addition, the backbone groups of K24, G56, W57, S58, G97, F99, P117 residues of TAX1BP1 form eight hydrogen bonds with the side chains of S37, H38, Y45, R51, and K53 from the paired NAP1 dimer (Fig 6D). Furthermore, two charge-charge interactions formed between the side chains of E98, R121 from TAX1BP1 and K49, E56 of NAP1 further contribute to the NAP1/TBK1 complex formation (Fig 6D).

### NDP52 and TAX1BP1 share a general binding mode to interact with NAP1 and SINTBAD

Detailed structural and sequence comparison analyses of the NAP1/NDP52 and NAP1/TAX1BP1 complexes revealed that the NAP1 dimer employs almost the same residues to interact with NDP52 and TAX1BP1, except for the L52 and K53 residues that are only important for binding to TAX1BP1 (Figs 4A, 6D and **EV1A**), and the corresponding residues involved in the interaction with NAP1 of NDP52 and TAX1BP1 are also highly similar (Figs 4A, 6A and D). Interestingly, like NAP1, another TBK1-binding adaptor, SINTBAD, was also reported to associate with NDP52 through its N-terminal coiled-coil region (Ryzhakov & Randow, 2007; Thurston et al, 2009). Careful sequence alignment analysis of the NDP52-binding regions in these two adaptor proteins revealed that most key residues of NAP1 that are critical for homo-dimer formation or interaction with NDP52 could be also found in SINTBAD (Fig 6E). Accordingly, based on these structural and sequence observations, we concluded that the autophagy receptor NDP52 and TAX1BP1 share a similar binding model to interact with NAP1 and SINTBAD, thereby forming distinct complexes, which are likely responsible for different selective autophagy processes.

### Evaluations of the TBK1-mediated phosphorylation sites in the SKICH domains of NDP52 and TAX1BP1 on the formation of NDP52/NAP1 or TAX1BP1/NAP1 complex

Recent studies reported that the TBK1 kinase can directly or indirectly mediate the phosphorylation of the T39 and S120 residues of NDP52 as well as the S25 residue of TAX1BP1, all of which are within the SKICH region of these two proteins (Richter et al, 2016). In our determined NDP52/NAP1 complex structure, the T39 residue of NDP52 is located in the β2-strand of the SKICH domain, and is far away from the NAP1-binding site (**Fig EV8A**), while the side chain of NDP52 S120 residue is buried in the structural core of the SKICH domain by forming two strong hydrogen bonds with the backbone amide of F107 and the backbone carboxyl group of I121, respectively (**Fig EV8A**). Apparently, the phosphorylation of NDP52 T39 residue is impossible to affect the interaction between NDP52 and NAP1, and the S120 residue is unlikely to be phosphorylated due to its crucial structural roles. Consistent with these structural observations, we were unable to get soluble recombinant protein for the phosphomimetic S120E mutant of NDP52(1-126), and further ITC-based measurement revealed that the phosphomimetic T39E mutant of NDP52(1-126) has a similar binding ability to NAP1(33-75) as that of the wild type protein (**Figs EV8A** and **1B**). Interestingly, the TAX1BP1 S25 residue is located in the loop connecting β1 and β2 strands, and is adjacent to the NAP1-binding interface in the TAX1BP1/NAP1 complex (**Fig EV8C**). Importantly, the side chain of S25 forms two hydrogen bonds with the side chain of TAX1BP1 R121 residue, which is further coupled with the negatively-charged E56 residue of NAP1 through a charge-charger interaction (**Fig EV8C**). Thus, once the S25 residue of TAX1BP1 was phosphorylated, the negatively charged phosphate group might disturb the interaction between TAX1BP1 and NAP1. In line with our structural analysis, further ITC-based measurement revealed that the phosphomimetic S25E mutant of TAX1BP1(1-121) showed a much weaker binding to NAP1(33-75) than that of the wild type TAX1BP1 protein (**Figs EV8D** and **5C**). Thus, TBK1 may mediate the phosphorylation of TAX1BP1 S25 residue to regulate the interaction between TAX1BP1 and NAP1.

## Discussion

In this study, we revealed that the N-terminal coiled-coil region of NAP1 forms a homo-dimer, and uses a similar binding mode and almost the identical key residues to associate with the SKICH regions of NDP52 and TAX1BP1, forming two unique hetero-tetramer complexes. Interestingly, although our analytical ultracentrifugation assay showed that the NAP1(33-75) dimer could form two stoichiometric types of complex with the monomeric NDP52(10-126) fragment under a un-saturated condition in solution (Fig 1G), further analyses showed that the dimeric form of NDP52 such as the NDP52(1-316) fragment, which includes the central coiled-coil region of NDP52, only forms a stable hetero-tetramer with the NAP1(33-75) dimer (**Fig EV9**). Therefore, the full length NDP52 and TAX1BP1, both of which were reported to form a dimer via their middle coiled-coil regions (Ling & Goeddel, 2000; Sternsdorf et al, 1997; Xie et al, 2015), are predicated to form hetero-tetrameric complexes when binding to NAP1. In addition, based on structural and sequence analyses, we inferred that the two TBK1-binding adaptors, NAP1 and SINTBAD, likely share a similar binding model to interact with NDP52 and TAX1BP1. In the future, it will be interesting to know the relationship between NAP1 and SINTBAD in binding to NDP52 and TAX1BP1, and whether NAP1 and SINTBAD play redundant or different roles in bridging TBK1 kinase to NDP52 and TAX1BP1 during different selective autophagy processes.

So far, the SKICH domain is identified in many different proteins including NDP52, TAX1BP1, CALCOCO1, SKIP and PIPP. However, due to lacking of detailed structure characterization, the precise working mode of SKICH domain is still elusive. Therefore, the structures of the NDP52/NAP1 and TAX1BP1/NAP1 complexes determined in this work provide the first atomic picture showing how a SKICH domain functions as a protein-protein interaction module to interact with its binding partners. Interestingly, although both SHICH domain of NDP52 and TAX1BP1 can bind to NAP1, detailed sequence and structural analyses revealed that the NAP1-binding residues are not conserved in the SKICH domains of CALCOCO1, SKIP and PIPP (**Fig EV10A** and **B**). Therefore, rather than binding to NAP1, the SKICH domains of CALCOCO1, SKIP and PIPP are likely to interact with other unknown proteins. Definitely, more work is required to clarify the functions of those SKICH domains.

In contrast to autophagy receptor Optineurin, which may directly bind to TBK1 as demonstrated by our previous study (Li et al, 2016), NDP52 and TAX1BP1 rely on NAP1 and/or SINTBAD to associate with TBK1. Previous studies indicated that TBK1, together with NDP52, TAX1BP1 and Optineurin, was recruited to the ubiquitin-decorated damaged mitochondria in the depolarization-dependent mitophagy as well as the invading pathogen in xenophagy, and cooperated with those autophagy receptors in selective autophagy (Heo et al, 2015; Lazarou et al, 2015; Richter et al, 2016; Thurston et al, 2016; Thurston et al, 2009). Importantly, TBK1 can directly phosphorylate Optineurin to regulate its interactions with Atg8 family proteins and ubiquitin proteins (Heo et al, 2015; Li et al, 2018; Richter et al, 2016; Wild et al, 2011). Strikingly, although TBK1 can also directly or indirectly phosphorylate NDP52 and TAX1BP1 (Heo et al, 2015; Richter et al, 2016), the precise downstream effects induced by TBK1-mediated phosphorylation of NDP52 and TAX1BP1 in selective autophagy are still unknown. In this study, we investigated the currently known phosphorylation sites mediated by TBK1 in the SKICH domains of NDP52 and TAX1BP1. Our data showed that the phosphorylation of NDP52 T39 residue is unable to affect the interaction between NDP52 and NAP1 but instead may alter the ability of NDP52 to interact with other unknown partners, and the S120 residue of NDP52 is unlikely to be phosphorylated as it is tightly packed in the structural core of the SKICH domain (**Fig EV8A** and **B**). Interestingly, the phosphorylation of TAX1BP1 S25 residue may reduce the interaction between TAX1BP1 and NAP1 (**Fig EV8C** and **D**), thereby likely to provide a negative feedback loop in TAX1BP1-mediated selective autophagy processes. Consistent with this notion, a recent study demonstrated that TBK1 could not promote the recruitment of TAX1BP1 to damaged mitochondria in the depolarization-dependent mitophagy, in contrast to NDP52 and Optineurin (Heo et al, 2015). Unfortunately, due to technological limitations, we were unable to evaluate the down-stream functional effects induced by this S25-phosphorylation of TAX1BP1 *in vivo*. Moreover, the activation of TBK1 in response to mitochondrial depolarization in mitophagy was proved to involve NDP52, but not TAX1BP1 (Heo et al, 2015), however, the exact underlying mechanism is still poorly understood. Therefore, further studies are required to elucidate the detailed relationships between TBK1 and two autophagy receptors, NDP52 and TAX1BP1.

In summary, we proposed a model depicting the recruitments of TBK1 to NDP52 and TAX1BP1 mediated by NAP1 as well as the potential regulations of NDP52 and TAX1BP1 by TBK1 in mitophagy and/or xenophagy (**Fig EV11**). In this model, NDP52 and TAX1BP1 both formed dimers through their coiled-coil regions, and separately associated with the NAP1 dimer mediated by the unique interactions between their SKICH domains and the N-terminal coiled-coil domain of NAP1 (**Fig EV11**). Meanwhile, the C-terminal TBD domains of the NAP1 dimer could further interact with the CTD domain of TBK1 dimer, thereby to form two hetero-hexamers (**Fig EV11**). Then, the NDP52/NAP1/TBK1 and TAX1BP1/NAP1/TBK1 hetero-hexameric complexes were recruited to the ubiquitin-decorated mitochondria and/or pathogen through the C-terminal ubiquitin-binding ZF domains of NDP52 and TAX1BP1, respectively (**Fig EV11**). Finally, after activation of TBK1, the activated TBK1 molecules could directly or indirectly phosphorylate NDP52 and TAX1BP1 to regulate their binding abilities to NAP1 (as for TAX1BP1) and/or other unknown binding partners.

## Materials and Methods

### Protein expression and purification

Different DNA fragments encoding human NDP52, NAP1, TAX1BP1 and other related DNA fragments were amplified by PCR from the full-length human cDNA, respectively. All these fragments were either cloned into the pET-32M vector (a modified version of pET32a vector containing a N-terminal Trx-tag and His_6_-tag) or the pET-GST vector (a modified version of pET32a vector containing a N-terminal GST-tag) for recombinant protein expressions. For the fluorescence imaging experiment or co-immunoprecipitation assay, the full-length NDP52 and NAP1 DNA fragments were cloned into pmCherry-C1 and pEGFP-C1 vectors, respectively. All the point mutations of NDP52 and NAP1 or other relevant mutations used in this study were created using the standard PCR-based mutagenesis method, further checked by PCR screen using 2×Taq Master Mix (Vazyme Biotech Co., Ltd.) enzyme and confirmed by DNA sequencing.

Recombinant proteins were expressed in BL21 (DE3) *E. coli* cells induced by 100 μM IPTG at 16°C. The bacterial cell pellets were re-suspended in the binding buffer (50 mM Tris, 500 mM NaCl, 5 mM imidazole at pH 7.9), and then lysed by the FB-110XNANO homogenizer machine (Shanghai Litu Machinery Equipment Engineering Co., Ltd.). Then the lysis was centrifuged at 35000 g for 30 minutes to remove the debris. His_6_-tagged proteins were purified by Ni^2+^-NTA agarose (GE Healthcare) affinity chromatography, while GST-tagged proteins were purified by glutathione sepharose 4B (GE Healthcare) affinity chromatography. Each recombinant protein was further purified by size-exclusion chromatography or mono-Q ion-exchange chromatography. The N-terminal tag of each recombinant protein was cleaved by 3C protease and further removed by size-exclusion chromatography. Uniformly ^15^N- or ^15^N/^13^C-labeled proteins were prepared by growing bacteria in M9 minimal medium using ^15^NH_4_Cl (Cambridge Isotope Laboratories Inc., NLM-467) as the sole nitrogen source or ^15^NH_4_Cl and ^13^C_6_-glucose (Cambridge Isotope Laboratories Inc., CLM-1396) as the sole nitrogen and carbon sources, respectively.

### Isothermal titration calorimetry assay

ITC measurements were carried out on a MicroCal PEAQ-ITC calorimeter or an automated system (Malvern) at 25 °C. All protein samples were in the same buffer containing 50 mM Tris (pH 7.5), 100 mM NaCl and 1 mM DTT. The titration processes were performed by injecting 40 μl aliquots of the syringe sample into the cell sample at time intervals of 2 minutes to ensure that the titration peak returned to the baseline. The titration data were analyzed using the Malvern MicroCal PEAQ-ITC analysis program.

### NMR spectroscopy

The stable isotope labeled protein samples for NMR studies were concentrated to ∼0.1 mM for titration experiments and ∼0.6 mM for backbone resonance assignment experiments in 50 mM potassium phosphate buffer containing 50 mM NaCl, and 1 mM DTT at pH 6.5. NMR spectra were acquired at 25°C on an Agilent 800 MHz spectrometer equipped with an actively z gradient shielded triple resonance cryogenic probe. Backbone resonance assignments were achieved using a suite of heteronuclear correlation experiments including HNCO, HNCA, CA(CO)NH, HNCACB and CBCA(CO)NH using ^15^N/^13^C-labeled protein samples together with a 3D ^15^N-seperated NOESY (Bax & Grzesiek, 1993).

### Analytical gel filtration chromatography

Analytical gel filtration chromatography was carried out on an AKTA FPLC system (GE Healthcare). Protein samples were loaded on to a Superose 12 10/300 GL column (GE Healthcare) equilibrated with a buffer containing 20 mM Tris-HCl (pH 7.5), 100 mM NaCl and 1 mM DTT.

### Analytical Ultracentrifugation

Sedimentation velocity experiments were performed on a Beckman XL-I analytical ultracentrifuge equipped with an eight-cell rotor under 42000rpm at 20°C. The partial specific volume of different protein samples and the buffer density were calculated using the program SEDNTERP (http://www.rasmb.bbri.org/). The final sedimentation velocity data were analyzed and fitted to a continuous sedimentation coefficient distribution model using the program SEDFIT (Schuck, 2000). The fitting results are further output to the Origin 9.0 software and aligned with each other.

### Protein crystallization and structural elucidation

Crystals of NDP52(10-126), NDP52(10-126)/NAP1(33-75) complex and TAX1BP1(1-121)/NAP1(33-75) complex were obtained using the sitting-drop vapor-diffusion method at 16 °C. Specifically, the freshly purified NDP52(10-126) protein (20 or 10 mg/ml in 20 mM Tris-HCl, 100 mM NaCl, 1 mM DTT, 1 mM EDTA at pH 7.5) was mixed with equal volume of reservoir solution containing 0.2 M potassium sulfate (pH 6.7), 20% (w/v) polyethylene glycol 3350. While, crystals of NDP52(10-126)/NAP1(33-75) complex (2.2 or 1.4 mg/ml in 50 mM Tris-HCl, 100 mM NaCl, 1 mM DTT, 1 mM EDTA at pH 7.5), and TAX1BP1(1-121)/NAP1(33-75) complex (20 or 10 mg/ml in 50 mM Tris-HCl, 100 mM NaCl, 1 mM DTT, 1 mM EDTA at pH 7.5) were grown from 0.2 M sodium malonate buffer at pH 7.0, 20% (w/v) polyethylene glycol 3350, and 0.1 M HEPES buffer at pH 7.5, 4% (w/v) polyethylene glycol 8000, respectively. Before diffraction experiments, appropriate glycerol was added as the cryo-protectant. X-ray data sets were collected at the beamline BL17U1 or BL19U1 of the Shanghai Synchrotron Radiation Facility. The diffraction data were processed and scaled using HKL2000 (Otwinowski & Minor, 1997).

The phase problem of NDP52(10-126) was solved by the molecular replacement method using the modified structure of NDP52(21-141) (PDB ID: 3VVV) with PHASER (Storoni et al, 2004). While, the phase problems of NDP52(10-126)/NAP1(33-75) complex and TAX1BP1(1-121)/NAP1(33-75) complex were solved by the molecular replacement method using our determined structures of NDP52(10-126) and NDP52(10-126)/NAP1(33-75) complex, respectively. All initial structural models were rebuilt manually using COOT (Emsley & Cowtan, 2004), and then refined using REFMAC (Murshudov et al, 1997), or PHENIX (Adams et al, 2002). The qualities of the final model were validated by MolProbity (Davis et al, 2007). The final refinement statistics of solved structures in this study were listed in Table EV1. All the structural diagrams were prepared using the program PyMOL (http://www.pymol.org/).

### Co-immunoprecipitation assay

HEK293 T cells transiently expressing proteins were harvested, washed with PBS buffer, and lysed for 1 hour at 4°C in lysis buffer containing 50 mM Tris-HCl (pH 7.8), 50 mM NaCl, 0.4% NP-40, 0.5 mM PMSF and protease inhibitor cocktail (AMRESCO). Lysates were centrifuged, and then supernatants were incubated with appropriate antibody pretreated rProtein G Agarose (Invitrogen) for 3 hours under rotation at 4°C. Precipitated proteins were washed with lysis buffer for 5 times, and then collected by brief centrifugation. Subsequently, the precipitated proteins were resolved in SDS-PAGE gel and detected by immune-blotting using a chemical luminescence-based detection method.

### Cell culture, transfection and fluorescence imaging

HeLa cells were cultured in Dulbecco’s modified Eagle’s medium (DMEM, Invitrogen) supplemented with 10% fetal bovine serum (FBS, Invitrogen). Co-transfections of GFP-NAP1 and mCherry-NDP52 or related mutant plasmids were performed with Lipofectamine 2000 (Invitrogen) according to the manufacturer’s instructions. After 48 hours, cells were fixed with 4% paraformaldehyde and punched with 0.2% Triton X-100/PBS, and the nuclei were visualized by staining with DAPI. The cell images were captured and analyzed using the TCS SP5 microscope equipped with LAS X software (Leica, Inc.). Particularly, the Pearson’s correlation of co-localization was performed using the LAS X software based on a randomly selected region that roughly contains one co-transfected HeLa cell. The statistical data represent mean ± SEM of >30 analyzed cells (selected regions) from two independent experiments. The unpaired Student t-test analysis was used to define a statistically significant difference.

### Coordinates

The coordinates and structure factors of the NDP52(10-126), NDP52(10-126)/NAP1(33-75) complex, and TAX1BP1(1-121)/NAP1(33-75) complex have been deposited in the Protein Data Bank under the accession code 5Z7A, 5Z7L and 5Z7G, respectively.

## Acknowledgements

We thank NCPSS BL19U1 and SSRF BL17U1 for X-ray beam time, Dr. Jianchao Li for help in the X-ray diffraction data collection, Prof. Jiahuai Han for the full length NDP52, TAX1BP1 cDNA, Dr. Chengjiang Gao for the NAP1 plasmid and Prof. Mingjie Zhang for the critical reading of the manuscript. This work was supported by grants from the National Natural Science Foundation of China (31470749, 21621002), the National Key R&D Program of China (2016YFA0501903), the Science and Technology Commission of Shanghai Municipality (15JC1400400), the Strategic Priority Research Program of the Chinese Academy of Sciences (XDB20000000), a “Thousand Talents Program” young investigator award, the start-up fund from State Key Laboratory of Bioorganic and Natural Products Chemistry and Chinese Academy of Sciences.

## Author contributions

T.F, Y.W. and L.P. designed the experiments. T.F. and X.X performed protein purifications, protein interaction assays, protein crystallizations and x-ray diffraction data collections. T.F. and L.P. determined the crystal structures. T.F. and Y.W. performed all the cell-based assays. J.L., X.X., Y.W., and S.H. helped in the data analysis. T.F. and L.P. analyzed the data and wrote the manuscript. L.P. supervised the project.

## References

Adams PD, Grosse-Kunstleve RW, Hung LW, Ioerger TR, McCoy AJ, Moriarty NW, Read RJ, Sacchettini JC, Sauter NK, Terwilliger TC (2002) PHENIX: building new software for automated crystallographic structure determination. Acta Crystallogr D 58: 1948–1954

Bax A, Grzesiek S (1993) Methodological Advances in Protein Nmr. Accounts Chem Res 26: 131–138

Davis IW, Leaver-Fay A, Chen VB, Block JN, Kapral GJ, Wang X, Murray LW, Arendall WB, Snoeyink J, Richardson JS, Richardson DC (2007) MolProbity: all-atom contacts and structure validation for proteins and nucleic acids. Nucleic Acids Res 35: W375–W383

Ellinghaus D, Zhang H, Zeissig S, Lipinski S, Till A, Jiang T, Stade B, Bromberg Y, Ellinghaus E, Keller A, Rivas MA, Skieceviciene J, Doncheva NT, Liu X, Liu Q, Jiang F, Forster M, Mayr G, Albrecht M, Hasler R, Boehm BO, Goodall J, Berzuini CR, Lee J, Andersen V, Vogel U, Kupcinskas L, Kayser M, Krawczak M, Nikolaus S, Weersma RK, Ponsioen CY, Sans M, Wijmenga C, Strachan DP, McArdle WL, Vermeire S, Rutgeerts P, Sanderson JD, Mathew CG, Vatn MH, Wang J, Nothen MM, Duerr RH, Buning C, Brand S, Glas J, Winkelmann J, Illig T, Latiano A, Annese V, Halfvarson J, D’Amato M, Daly MJ, Nothnagel M, Karlsen TH, Subramani S, Rosenstiel P, Schreiber S, Parkes M, Franke A (2013) Association between variants of PRDM1 and NDP52 and Crohn’s disease, based on exome sequencing and functional studies. Gastroenterology 145: 339–347

Emsley P, Cowtan K (2004) Coot: model-building tools for molecular graphics. Acta Crystallogr D 60: 2126–2132

Feng YC, He D, Yao ZY, Klionsky DJ (2014) The machinery of macroautophagy. Cell Res 24: 24–41

Gibbings D, Mostowy S, Jay F, Schwab Y, Cossart P, Voinnet O (2012) Selective autophagy degrades DICER and AGO2 and regulates miRNA activity. Nature cell biology 14: 1314–1321

Gomes LC, Dikic I (2014) Autophagy in antimicrobial immunity. Mol Cell 54: 224–233

Guo H, Chitiprolu M, Gagnon D, Meng L, Perez-Iratxeta C, Lagace D, Gibbings D (2014) Autophagy supports genomic stability by degrading retrotransposon RNA. Nat Commun 5: 5276

Gurung R, Tan A, Ooms LM, McGrath MJ, Huysmans RD, Munday AD, Prescott M, Whisstock JC, Mitchell CA (2003) Identification of a novel domain in two mammalian inositol-polyphosphate 5-phosphatases that mediates membrane ruffle localization. The inositol 5-phosphatase skip localizes to the endoplasmic reticulum and translocates to membrane ruffles following epidermal growth factor stimulation. The Journal of biological chemistry 278: 11376–11385

Heo JM, Ordureau A, Paulo JA, Rinehart J, Harper JW (2015) The PINK1-PARKIN Mitochondrial Ubiquitylation Pathway Drives a Program of OPTN/NDP52 Recruitment and TBK1 Activation to Promote Mitophagy. Mol Cell 60: 7–20

Holm L, Rosenstrom P (2010) Dali server: conservation mapping in 3D. Nucleic Acids Res 38: W545–549

Jiang P, Mizushima N (2014) Autophagy and human diseases. Cell Res 24: 69–79

Jin S, Tian S, Luo M, Xie W, Liu T, Duan T, Wu Y, Cui J (2017) Tetherin Suppresses Type I Interferon Signaling by Targeting MAVS for NDP52-Mediated Selective Autophagic Degradation in Human Cells. Mol Cell 68: 308–322 e304

Johansen T, Lamark T (2011) Selective autophagy mediated by autophagic adapter proteins. Autophagy 7: 279–296

Kim BW, Hong SB, Kim JH, Kwon do H, Song HK (2013) Structural basis for recognition of autophagic receptor NDP52 by the sugar receptor galectin-8. Nat Commun 4: 1613

Klionsky DJ, Emr SD (2000) Cell biology - Autophagy as a regulated pathway of cellular degradation. Science 290: 1717–1721

Lazarou M, Sliter DA, Kane LA, Sarraf SA, Wang C, Burman JL, Sideris DP, Fogel AI, Youle RJ (2015) The ubiquitin kinase PINK1 recruits autophagy receptors to induce mitophagy. Nature 524: 309–314

Li F, Xu D, Wang Y, Zhou Z, Liu J, Hu S, Gong Y, Yuan J, Pan L (2018) Structural insights into the ubiquitin recognition by OPTN (optineurin) and its regulation by TBK1-mediated phosphorylation. Autophagy: 1–14

Li FX, Xie XQ, Wang YL, Liu JP, Cheng XF, Guo YJ, Gong YK, Hu SC, Pan LF (2016) Structural insights into the interaction and disease mechanism of neurodegenerative disease-associated optineurin and TBK1 proteins. Nat Commun **7**

Li S, Wandel MP, Li F, Liu Z, He C, Wu J, Shi Y, Randow F (2013) Sterical hindrance promotes selectivity of the autophagy cargo receptor NDP52 for the danger receptor galectin-8 in antibacterial autophagy. Sci Signal 6: ra9

Ling L, Goeddel DV (2000) T6BP, a TRAF6-interacting protein involved in IL-1 signaling. Proceedings of the National Academy of Sciences of the United States of America 97: 9567–9572

Matsumoto G, Wada K, Okuno M, Kurosawa M, Nukina N (2011) Serine 403 phosphorylation of p62/SQSTM1 regulates selective autophagic clearance of ubiquitinated proteins. Mol Cell 44: 279–289

Mizushima N (2007) Autophagy: process and function. Genes & development 21: 2861–2873

Moore AS, Holzbaur EL (2016) Dynamic recruitment and activation of ALS-associated TBK1 with its target optineurin are required for efficient mitophagy. Proceedings of the National Academy of Sciences of the United States of America 113: E3349–3358

Murshudov GN, Vagin AA, Dodson EJ (1997) Refinement of macromolecular structures by the maximum-likelihood method. Acta Crystallogr D 53: 240–255

Nakatogawa H, Suzuki K, Kamada Y, Ohsumi Y (2009) Dynamics and diversity in autophagy mechanisms: lessons from yeast. Nat Rev Mol Cell Bio 10: 458–467

Newman AC, Scholefield CL, Kemp AJ, Newman M, McIver EG, Kamal A, Wilkinson S (2012) TBK1 kinase addiction in lung cancer cells is mediated via autophagy of Tax1bp1/Ndp52 and non-canonical NF-kappaB signalling. PloS one 7: e50672

Otwinowski Z, Minor W (1997) Processing of X-ray diffraction data collected in oscillation mode. Method Enzymol 276: 307–326

Randow F, Youle RJ (2014) Self and Nonself: How Autophagy Targets Mitochondria and Bacteria. Cell host & microbe 15: 404–412

Richter B, Sliter DA, Herhaus L, Stolz A, Wang C, Beli P, Zaffagnini G, Wild P, Martens S, Wagner SA, Youle RJ, Dikic I (2016) Phosphorylation of OPTN by TBK1 enhances its binding to Ub chains and promotes selective autophagy of damaged mitochondria. Proceedings of the National Academy of Sciences of the United States of America 113: 4039–4044

Rogov V, Dotsch V, Johansen T, Kirkin V (2014) Interactions between Autophagy Receptors and Ubiquitin-like Proteins Form the Molecular Basis for Selective Autophagy. Mol Cell 53: 167–178

Ryzhakov G, Randow F (2007) SINTBAD, a novel component of innate antiviral immunity, shares a TBK1-binding domain with NAP1 and TANK. Embo J 26: 3180–3190

Schuck P (2000) Size-distribution analysis of macromolecules by sedimentation velocity ultracentrifugation and lamm equation modeling. Biophys J 78: 1606–1619

Sternsdorf T, Jensen K, Zuchner D, Will H (1997) Cellular localization, expression, and structure of the nuclear dot protein 52. The Journal of cell biology 138: 435–448

Stolz A, Ernst A, Dikic I (2014) Cargo recognition and trafficking in selective autophagy. Nature cell biology 16: 495–501

Storoni LC, McCoy AJ, Read RJ (2004) Likelihood-enhanced fast rotation functions. Acta Crystallogr D 60: 432–438

Thurston TL, Boyle KB, Allen M, Ravenhill BJ, Karpiyevich M, Bloor S, Kaul A, Noad J, Foeglein A, Matthews SA, Komander D, Bycroft M, Randow F (2016) Recruitment of TBK1 to cytosol-invading Salmonella induces WIPI2-dependent antibacterial autophagy. Embo J 35: 1779–1792

Thurston TL, Ryzhakov G, Bloor S, von Muhlinen N, Randow F (2009) The TBK1 adaptor and autophagy receptor NDP52 restricts the proliferation of ubiquitin-coated bacteria. Nat Immunol 10: 1215–1221

Thurston TL, Wandel MP, von Muhlinen N, Foeglein A, Randow F (2012) Galectin 8 targets damaged vesicles for autophagy to defend cells against bacterial invasion. Nature 482: 414–418

Tumbarello DA, Manna PT, Allen M, Bycroft M, Arden SD, Kendrick-Jones J, Buss F (2015) The Autophagy Receptor TAX1BP1 and the Molecular Motor Myosin VI Are Required for Clearance of Salmonella Typhimurium by Autophagy. PLoS pathogens 11: e1005174

Tumbarello DA, Waxse BJ, Arden SD, Bright NA, Kendrick-Jones J, Buss F (2012) Autophagy receptors link myosin VI to autophagosomes to mediate Tom1-dependent autophagosome maturation and fusion with the lysosome. Nature cell biology 14: 1024–1035

von Muhlinen N, Akutsu M, Ravenhill BJ, Foeglein A, Bloor S, Rutherford TJ, Freund SM, Komander D, Randow F (2012) LC3C, bound selectively by a noncanonical LIR motif in NDP52, is required for antibacterial autophagy. Mol Cell 48: 329–342

Wild P, Farhan H, McEwan DG, Wagner S, Rogov VV, Brady NR, Richter B, Korac J, Waidmann O, Choudhary C, Dotsch V, Bumann D, Dikic I (2011) Phosphorylation of the autophagy receptor optineurin restricts Salmonella growth. Science 333: 228–233

Xie X, Li F, Wang Y, Wang Y, Lin Z, Cheng X, Liu J, Chen C, Pan L (2015) Molecular basis of ubiquitin recognition by the autophagy receptor CALCOCO2. Autophagy 11: 1775–1789

